# Interrelations between delta waves, spindles and slow oscillations in human NREM sleep and their functional role in memory

**DOI:** 10.1101/2021.09.03.458607

**Authors:** Larissa N. Wüst, Daria Antonenko, Robert Malinowski, Liliia Khakimova, Ulrike Grittner, Klaus Obermayer, Julia Ladenbauer, Agnes Flöel

## Abstract

Certain neurophysiological characteristics of sleep, in particular slow oscillations (SO), sleep spindles, and their temporal coupling, have been well characterized and associated with human memory formation. Delta waves, which are somewhat higher in frequency and lower in amplitude compared to SO, have only recently been found to play a critical role in memory processing of rodents, through a competitive interplay between SO-spindle and delta-spindle coupling. However, human studies that comprehensively address delta waves, their interactions with spindles and SOs as well as their functional role for memory are still lacking.

Electroencephalographic data were acquired across three naps of 33 healthy older human participants (17 female) to investigate delta-spindle coupling and the interplay between delta and SO-related activity. Additionally, we determined intra-individual stability of coupling measures and their potential link to the ability to form novel memories.

Our results revealed weaker delta-spindle compared to SO-spindle coupling. Contrary to our initial hypothesis, we found that increased delta activity was accompanied by stronger SO-spindle coupling. Moreover, we identified the ratio between SO- and delta-nested spindles as the sleep parameter that predicted ability to form novel memories best.

Our study suggests that SOs, delta waves and sleep spindles should be jointly considered when aiming to link sleep physiology and memory formation in aging.

**Significance statement:** Interrelations between delta waves, slow oscillations and sleep spindles have recently been causally linked to the balance between consolidation and forgetting in rats using optogenetics. In humans, SO-spindle coupling has been intensively investigated, but delta waves and their interaction with spindles were only studied jointly as SWA. Here we delineate the coupling of delta waves to spindles, investigate interactions of delta- and SO-related activity and investigate their role for the ability to form novel memories in healthy older individuals. Our results show differences as well as dependencies between SO- and delta-related activities including spindle coupling. Further, our results indicate that the ratio of SO- to delta-nested spindles might be the most informative sleep parameter for memory formation of older adults.

## Introduction

Specific brain oscillations during human non-rapid eye movement (NREM) sleep, most importantly thalamo-cortical spindles (12-15 Hz) and cortical slow wave activity (SWA) which can be divided into slow oscillations (SO, 0.5-1 Hz) and delta waves (1-4 Hz) (Genzel et al., 2014; Navarrete et al., 2020), have been linked to successful memory consolidation, and thus, the ability to retain novel memories (Rasch and Born, 2013). The differentiation between slow waves can be made based on their origin, extent and amplitude (Siclari et al., 2014; Navarrete et al., 2020). While both SO and delta waves originate in cortico-cortical networks of primary and association cortices, the thalamus contributes to the generation of the SOs with larger amplitude (< 1 Hz) and mediates its large-scale synchronization (Lemieux et al., 2014).

In humans, the role of SOs for memory processes has been extensively investigated. As such, previous studies have shown increased SO activity after learning, correlating with retention performance after sleep (Gais and Born, 2004; Mölle et al., 2004; Mölle et al., 2009; Rasch and Born, 2013). There is also growing evidence that temporal coordination of SOs with spindles, rather than SO activity in isolation, is crucial for memory consolidation (Ladenbauer et al., 2017; Helfrich et al., 2018; Muehlroth et al., 2019). Moreover, SO-spindle coupling was shown to substantially vary between individuals, but to be stable intra-individually, indicating that this parameter may be a “fingerprint-like” general trait (Cox et al., 2018). This observation may have implications for individual memory abilities beyond sleep-related memory consolidation, namely the ability to form novel memories.

With regard to delta waves most studies have considered SO and delta activity jointly as SWA, thus precluding investigation of delta coupling behavior with spindles, as well as any attempt to delineate their distinct contribution to memory processes (Ngo and Born, 2019).

One recent rodent study suggested a causal role for delta waves and their interaction with spindles for memory (Kim et al., 2019). In this study, disturbance during delta wave up-states by means of optogenetic stimulation led to a relative increase in SO-nested spindles, enhanced reactivation strength in involved neuronal ensembles, and improved memory performance after sleep. Disturbance during SOs had the opposite effect, resulting in a relatively higher number of spindles nested to delta waves, reduced ensemble reactivations, and impaired performance, suggesting the following: First, there seems to be an opposing dependency between SO and delta wave generating systems; second, SO and delta waves may have competitive functional roles in memory (Ngo and Born, 2019). While SO-related spindles support memory consolidation, delta-related spindles seem to benefit forgetting, possibly to support downscaling processes during sleep to restore the brain’s capacity to learn (Kim et al., 2019). Interestingly, not SO-spindle coupling by itself, but the ratio between SO-nested spindles to delta-nested spindles was the sleep parameter most predictive for memory performance gains after sleep (Kim et al., 2019). This finding suggests that both SO-nested and delta-nested spindles may have to be taken into account, when relating sleep physiology to memory. However, delta-spindle coupling has not been investigated in humans yet, and evidence on its functional role is limited. Moreover, it is unknown whether there is a competitive interaction between SO and delta wave activities in humans. This information may be particularly valuable when evaluating interventions that are aimed to modulate memory abilities in healthy and pathological aging.

In the current study, we therefore first aimed to characterize delta-spindle coupling in healthy older adults, and to compare it to SO-spindle coupling. Second, we aimed to address the previously described competitive interplay between SO- and delta-related activities in natural human nap sleep by evaluating SO-spindle coupling during sleep periods with low compared to high delta activity. Third, we investigated stability of SO- and delta-spindle coupling measures over multiple nap sleep assessments to reveal whether these sleep parameters exhibit a trait-like character. Finally, we tested for associations between ability to form novel memories and SO- and delta-spindle coupling measures in healthy older participants, including the ratio between SO-nested spindles to delta-nested spindles.

## Materials and Methods

### Participants and experimental procedure

Thirty three healthy older adults (17 female, age range: 50 to 78 years) were included in the study. Exclusion criteria included history of severe untreated medical, neurological, and psychiatric diseases; subjective cognitive decline; sleep disorders; cognitive impairment; intake of medication acting primarily on the central nervous system (e.g., antipsychotics, antidepressants, benzodiazepines, or any type of over-the-counter sleep-inducing drugs like valerian); alcohol or substance abuse. Our choice to investigate healthy older individuals was based on the motivation of understanding and eventually improving sleep physiology and memory in aging. Data were collected within the scope of an extensive study involving several sets of transcranial direct current stimulation (tDCS) protocols, in addition to the sessions without tDCS that were used in this report (9 sleep sessions in total; 3 sleep sessions “without tDCS”: T1, T2, T3). Whereas T1 was always followed by T2 nap session, T3 was one of the 7 following sessions (randomized order; ClinicalTrials.gov Identifier: NCT04714879). The three naps were separated by ~ 5-6 weeks (mean ± SD time between T1 and T2: 31.2 ± 41.1 days; mean ± SD time between T2 and T3: 45.2 ± 32.1 days). A comprehensive neuropsychological testing was administered for assessment of cognitive status in each participant (Ladenbauer et al., 2016; Ladenbauer et al., 2017). Here, we used the verbal learning and memory test to probe memory performance (see below), which was acquired prior sleep on T1. In each session, participants arrived at the laboratory between 11:30 am and 2 pm (at a time suiting their daily routine to allow them to sleep). After preparation of polysomnographic recordings, participants were given the opportunity to sleep for max. 90 min. During the naps, EEG was recorded. The study was approved by the local ethics committee (Universitätsmedizin Greifswald) and conducted in accordance with the Declaration of Helsinki. Participants gave written informed consent prior to participation.

### Memory assessment

As part of the neuropsychological assessment, the German version of the Auditory Verbal Learning Test (AVLT) was performed by all participants to probe memory performance (Helmstaedter et al., 2001). Participants had to learn a list of 15 unrelated words within five immediate auditory presentations, each followed by an attempted recall (termed encoding trials). Subsequently a second 15-word interference list was presented, followed by retrieval of the initial list. Delayed recall and recognition were also tested after 30 minutes. Ability to form novel memories for each participant was indexed by the sum of recalled words from all five encoding trials (encoding performance) as well as by the difference between last encoding and first recall trial after interference (immediate recall performance) (Göder et al., 2013; Mikutta et al., 2019).

### EEG recordings

Polysomnographic data were recorded with a 32-channel Brain Products actiCAP electrode system positioned according to the 10-20 EEG system (Fp1, Fp2, AFz, Fz, F3, F4, F7, F8, FC1, FC2, FC5, FC6, Cz, C3, C4, T7, T8, CP1, CP2, CP5, CP6, Pz, P3, P4, P7, P8, O1, O2, reference: nose, ground: FCz) using the BrainAmp DC amplifier (Brain Products GmbH, Munich, Germany) with a sampling rate of 500 Hz. Horizontal and vertical electrooculogram (EOG) (using two electrodes in T1/T2, and four electrodes in T3) and the electromyogram (positioned at the chin) were recorded. Impedances were kept below 10 kOhm.

### EEG data analysis

EEG data was analyzed using BrainVision Analyzer 2.1 (Brain Products GmbH) and Matlab R2019b (The MathWorks, Inc.) with functions from the FieldTrip (Oostenveld et al., 2011) and CircStat toolboxes (Berens, 2009) as well as custom functions. Prior to sleep scoring and further analysis data were band-pass filtered (IIR filter, 0.032 - 35 Hz, notch: 50 Hz). To eliminate arousal and ocular artifacts, a semiautomatic detection procedure was applied including visual inspection of raw data (Ladenbauer et al., 2017) and an ocular correction procedure by means of independent component analysis (ICA). Sleep stages were scored based on standard criteria (Rechtschaffen and Kales, 1968). Only naps with a minimum duration of sleep stages 2, 3, and 4 of 15 min were included in the analysis, resulting in the following number of excluded naps: 3 in T1, 2 in T2, and 1 in T3. To characterize the temporal relationship between SOs and spindles, as well as delta waves and spindles, we performed time-frequency representations (TFR) and phase-amplitude coupling (PAC) analysis for frontal (Fz, FC1, FC2) and centroparietal (Cz, CP1, CP2) region-of-interests (ROI) (Ladenbauer et al., 2016; Ladenbauer et al., 2017). These were calculated on 30-s non-overlapping epochs of NREM sleep stage 2, 3 and 4 (sampling rate 500 Hz).

### SO, Delta wave, and spindle detection

Sleep oscillations of interest were detected using procedures previously applied in several studies (Mölle et al., 2002; Mölle et al., 2011; Staresina et al., 2015; Ladenbauer et al., 2017; Helfrich et al., 2018; Ladenbauer et al., 2021). For SO event detection, EEG-data were band-pass filtered at 0.16 to 1.25 Hz. All downward zero crossings were marked as potential start- and endpoints of discrete SO oscillations. The time between two successive downward zero crossings was determined and events were considered SOs for durations between 1 and 2 s, corresponding to a frequency of 0.5 to 1.0 Hz. For all selected events, the trough-to-peak amplitude was determined. The 25 % of events with the largest amplitudes were considered SOs and included into further analysis. For delta wave detection, EEG-data was band-pass filtered at 0.75 to 4.25 Hz. Time between successive downward zero crossings needed to be 0.25 to 1 s, corresponding to a frequency of 1 to 4 Hz for an event to be considered a delta wave. Only the 25 % of events with the highest amplitudes were considered delta waves and included into further analysis. For spindle detection, EEG-data was band-pass filtered at 12 to 15 Hz. Root mean square (RMS) values of the filtered signal were calculated with a moving average of 200 ms. Events with RMS amplitude values above an individual threshold determined as the 25 % of spindle events with the highest RMS amplitude values for a duration of 0.5 to 3 s were considered spindle events. Artifact-free segments of ±2.5 s around the event centers (SO or delta wave trough or spindle center) were extracted from the unfiltered signal for further analysis. To address the issue of spurious cross-frequency coupling, which can be caused by differences in oscillatory power (Aru et al., 2015), we performed a normalization of individual events to minimize amplitude differences prior to all subsequent analyses (Helfrich et al., 2018).

### Temporal relationships of SOs, Delta-waves, and spindles

#### TFR

For TFR analyses, detected SO and delta event segments were normalized using a z-score based on the mean and standard deviation calculated from the averaged event segment per participant and channel as established in previous studies (Helfrich et al., 2018; Ladenbauer et al., 2021). TFRs were calculated to display spectral densities of frequencies between 5 to 20 Hz in a time window of ±1.25 s around the SO trough. Therefore, spectral densities were calculated in 0.25 Hz steps using a sliding (10 ms steps) Hanning tapered window with a variable, frequency-dependent length. The window length always comprised a full number of five cycles to ensure reliable power estimates._ The baseline interval (−2.5 to −1.25 s) was used for a baseline correction of TFRs. The TFRs were then averaged across all segments and all electrodes of the respective ROI for each nap. The same procedure was applied to the delta event segments.

#### PAC

To compare SO and delta events in terms of their temporal interaction with spindles, we additionally analyzed their phase relationship, which allows a more precise characterization independent of their frequency. We determined cross-frequency coupling between SOs and spindles based on previous studies (Staresina et al., 2015; Ladenbauer et al., 2017; Helfrich et al., 2018; Ladenbauer et al., 2021), thus obtaining two parameters for each participant and nap: the SO phase of peak spindle power and the resultant vector length that represents the coupling strength between the two oscillations. We therefore filtered the normalized SO event segments in the SO frequency band (0.16 to 1.25 Hz) and extracted the instantaneous phase angle using the Hilbert transform. Subsequently, the same segments were filtered in the spindle frequency band (12-15 Hz) and the instantaneous amplitude was extracted from the Hilbert transform. We only considered the time range −2 to 2 s to avoid filter edge artifacts. In a next step, the maximum spindle amplitude and corresponding SO phase was extracted for each participant, nap, channel, and SO event. Finally, we determined the mean SO phase and corresponding resultant vector length across all events for each participant and nap using the CircStat toolbox (Berens, 2009) for MATLAB. The same procedures were used to determine delta-spindle PAC, using a filter of 0.75 to 4.25 Hz for the delta frequency band.

##### SO/delta nesting index

In addition to the two above described approaches, that characterize delta-spindle interactions and SO-spindle interactions in more detail, we calculated a sleep parameter that takes both SO-spindle and delta-spindle activity into account and was found to be the most predictive for memory consolidation in rats (Kim et al., 2019). Therefore, nested spindle events were defined according to previously used criteria (amplitude: 25 % highest amplitudes, duration: 0.5 to 3 s), with their center occurring within −0.5 to +0.5 s around the up-state of the SO or delta wave, respectively. If a spindle occurred within the nesting window of both, a SO and a delta wave, it was allocated to the event with the smallest time difference between the slow wave up-state and the spindle center. To determine SO-nesting, all spindles nested to SOs were counted. Delta-nesting was determined by counting the number of delta-nested spindles in each nap. Equivalent to Kim et al. (2019), we calculated the SO/delta nesting index for each nap by dividing the number of SO-nested spindles by the number of delta-nested spindles. Nesting analyses were calculated for each nap separately. To explore associations with memory performance, averages were calculated over all nap conditions for SO-spindle and delta-spindle coupling measures as well as for the SO/delta nesting index.

### Interdependence between delta activity and SO-related spindles

In light of recently published results on the interdependence between SO-related and delta-related activity in rats using optogenetic stimulation (Kim et al., 2019), we further addressed this issue in natural human sleep. We therefore compared SO-spindle coupling for low vs. high delta power segments. According to the findings in rats, SO-spindle coupling was expected to deviate depending on delta power. To this end, we determined delta power by applying a Hilbert transformation on the delta band-pass (1-4 Hz) filtered data. Delta power was averaged over the 30-s epoch and all electrodes of the respective ROI. A within-subject median split was applied to distinguish between epochs of low and high delta power for each nap (Iacobucci et al., 2015). TFR analysis for SO events and PAC analysis for SO-spindle associations were repeated for low and high delta power intervals.

### Statistical analysis

To test for event-locked power differences in the TFRs, we applied two-tailed paired-samples t-tests (group level statistics) to compare fast spindle power fluctuations in SOs versus delta waves, as well as in SO events of low versus high delta power intervals. Correction for multiple comparisons (−1.2 s to + 1.2 s, 5 to 20 Hz) was performed using a cluster-based permutation procedure as implemented in FieldTrip (Maris and Oostenveld, 2007; Staresina et al., 2015; Ladenbauer et al., 2017). V-tests (Berens, 2009) were used to characterize and compare SO-spindle and delta-spindle coupling phases separately for each nap. Given that phase is a circular metric, traditional analyses for repeated measures were not applicable. In line with previous studies (Staresina et al., 2015; Ladenbauer et al., 2017; Helfrich et al., 2019), we expected maximal spindle power around the SO up-state (i.e., around 0 deg), and compared whether this was also the case for delta-spindle coupling.

Statistical analyses for linear metrics were conducted with R (R Core Team, 2020). Linear mixed effect models (random intercept models) were computed for the dependent variables resultant vector length (coupling strength) and SO/delta nesting index. The repeated measurements (factor nap condition: T1, T2, T3) were level-one units nested in individuals (level-two units). Models included fixed effects for the oscillation (SO or delta wave), or delta power (low or high), respectively.

To test for intra-individual stability of relevant coupling measures across all nap conditions, intra-class correlation coefficients (ICC) were calculated for both SO-spindle and delta-spindle coupling strengths (resultant vector lengths) and the SO/delta nesting index, previously used to test trait-specific nature of variability in sleep/wake parameters (van Dongen et al., 2005; Tucker et al., 2007). Resulting estimates indicate the stability of the inter-individual differences across the three naps and were interpreted according to benchmark ranges: ‘slight’ (0.0–0.2), ‘fair’ (0.2–0.4), ‘moderate’ (0.4–0.6), ‘substantial’ (0.6–0.8), and ‘almost perfect’ (0.8–1.0) (Landis and Koch, 1977; Tucker et al., 2007). Pearson’s r correlation coefficients were then computed to test for a linear relationship between memory performance and the averaged SO-spindle, delta-spindle coupling strength or the averaged SO/delta nesting index over all nap conditions for each participant. No corrections for multiple comparisons were applied in this exploratory study. A two-sided significance level of α = 0.05 was used.

## Results

Participants slept on average for 81 min (mean ± SD in min: T1: 79.7 ± 15.2), T2: 81.1 ± 16.2, T3: 81.1 ± 13.0). Sleep architecture was comparable with previous reports in healthy older adults during daytime napping, with large proportion of lighter NREM sleep and only short to none SWS and REM sleep (mean ± SD in min: time spent in wake: 6.0 ± 7.5, sleep stage 1: 24.2 ± 15.4, sleep stage 2: 44.8 ± 16.8, sleep stage 3 and 4: 3.9 ± 6.5, REM: 1.3 ± 4.1) (Baran et al., 2016; Ladenbauer et al., 2021). As results for TFR and PAC analyses were similar between all nap conditions, we mainly report and depict results for T1, unless stated otherwise. As the same applies for the two ROIs, we depict results for the frontal ROI only.

Given that 1 Hz was the lower and upper bound of delta waves and SOs, respectively, we assessed how many events were included in both the delta and SO analyses and found about 6 events in each participant and nap. Since we consider this a negligible proportion of all SO (3 %) and delta events (0.5 %), we did not address this issue in the following analysis.

### SO-spindle coupling was stronger than delta-spindle coupling

TFRs over SO and delta events are shown in Figure 1A. As expected, averaged SO amplitude was higher in comparison to delta wave amplitude. Typical modulation of fast spindle power during SO events were found to be similar for delta events, with decreased power during the down-state, increased spindle power during the subsequent down-to-up transition and peaking around the up-state. While numerically, fast spindle power was higher around the up-states of delta waves than around the up-states of SOs, differences were only found for fast spindle power prior to the downstate as well as for the slow spindle frequency range (10-12 Hz).

**Figure 1.**
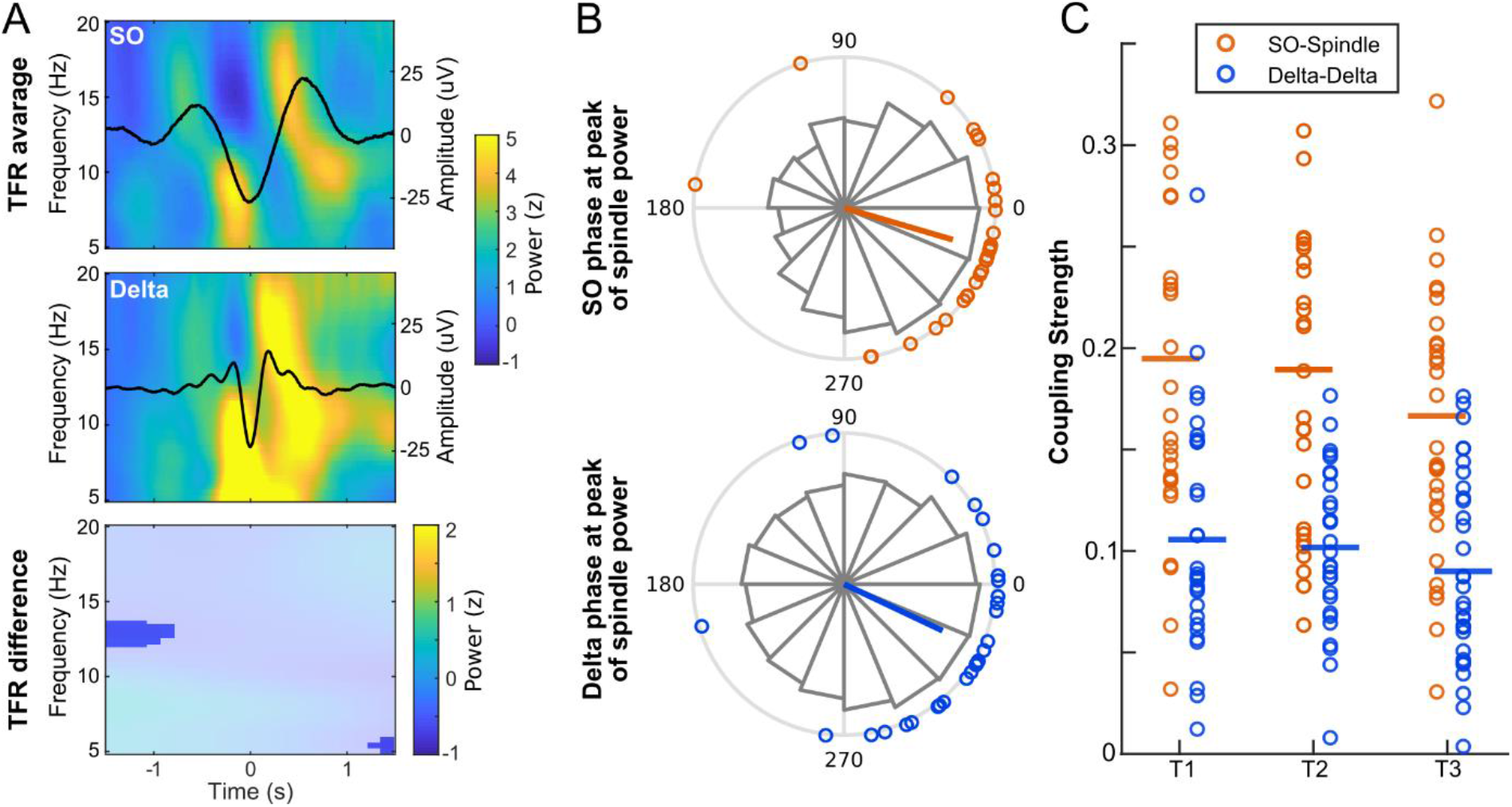
A: Average time-frequency representations (TFR) locked to the SO trough or delta trough (time 0), respectively. Shown are averaged power z-scores relative to pre-event baseline distributions. Black lines indicate averaged SO (top, n =175) or delta events (center, n = 1091) per participant. Bottom, Difference of these TFRs (SO-delta) masked by significance (p < 0.05, corrected for multiple comparison). B: SO-spindle and delta-spindle phase-amplitude coupling (PAC). Histogram indicates distribution of SO phase or delta phase angle at peak of spindle power across all SO or delta events, respectively. Circles at the circumference represent averages per participant and the colored line denotes the corresponding resulting vector length across all participants. C: Coupling strength (resultant vector length) for SO-spindle (orange) and delta-spindle coupling (blue) per participant. Circles represent individual participants and horizontal lines the average coupling strength across all participants for each nap condition (T1, T2, T3).

PAC for SO-spindle and delta-spindle coupling are shown in Figure 1B. Preferred phases of peak spindle power (i.e., mean phase of the SO or delta wave, respectively, at which the peak spindle power occurs) were comparable for SO-spindle (−16.32 ± 40.66 degrees, circular mean ± SD) and delta-spindle coupling (−25.18 ± 46.31 degrees), with spindle power maxima occurring in the late rising phase of the slow waves (albeit slightly more synchronous for SO). Concentrations around the SO and delta up-state (around 0 degree), were likewise found to be similar for SO-spindle (*V* = 20.83, *p* < .001) and delta-spindle coupling (*V* = 17.67, *p* < .001).

However, a substantial difference between SO-spindle and delta-spindle coupling emerged for coupling strength; i.e., resultant vector length. Due to the linearity of this metric, we were able to include data of all nap conditions by incorporating the factor nap condition in the model.

We found higher coupling strengths of spindles to SOs as compared to delta waves (beta: 0.084, 95% CI [0.069, 0.099], t(144.7) = 9.042, p < .001, R^2^ = .356; see **Fig. 1C**).

### SO-spindle coupling strength was lower in low compared to high delta power intervals

To test for interdependence between SO and delta related activity, separate TFR and PAC analyses of SO events were conducted for low and high delta power segments.

Time–frequency analysis revealed no substantial differences in fast spindle power between low and high delta power segments (**Fig. 2A**). Of note, the amplitude of the average SO event was smaller in low delta power intervals than in high delta power intervals, as was to be expected given the adjacent frequency range. Potential confounds that may have resulted from these amplitude differences in the following PAC analyses (Aru et al., 2015), however, were addressed by normalizing individual SO events (see methods).

**Figure 2.**
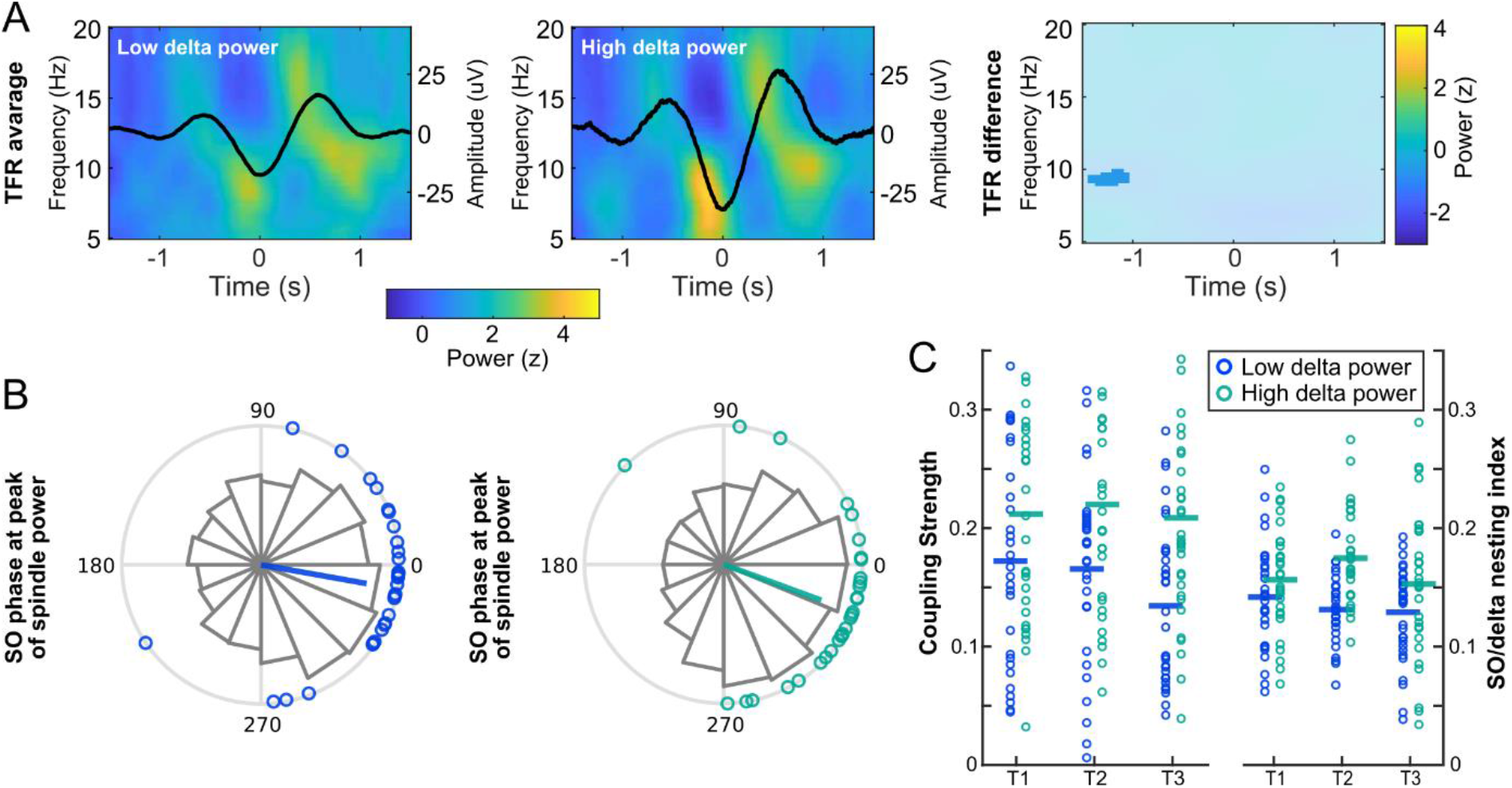
A: Average TFR for SO events in intervals of low (left, n = 83) and high (center, n = 93) delta power. Shown are averaged power z-scores relative to pre-event baseline distributions. Black lines indicate average SO events per participant. Difference of low – high delta power TFRs masked by significance (right, p < 0.05; corrected for multiple comparison). B: SO-spindle PAC for intervals of low (left) and high (right) delta power, respectively. Histogram indicates distribution of SO phase angle at peak of spindle power across all SO events. Circles at the circumference represent averages per participant and the colored line denotes the corresponding resulting vector length across all participants. C: Coupling strength and SO/delta nesting index for SO-spindle coupling per participant in intervals of low (blue) and high (cyan) delta power. SO/delta nesting index is the ratio of SO-nested spindles and delta nested spindles. Circles represent individual participants and horizontal lines the average coupling strength across all participants for each nap condition (T1, T2, T3).

PAC analyses addressing the SO-spindle phase relationships in these segments, revealed differences between low and high delta power segments. On the one hand, higher concentrations of spindle power maxima were found around the SO up-states for low delta power segments (across all SO events as well as for participant-averaged SO phases; see histogram and circles in **Fig. 2B**; Low: *V*=22.04, *p*<0.001; High: *V*=20.37, *p*<0.001), indicating more synchronous SO-spindle coupling in low compared to high delta power intervals. This was also underlined by the mean SO phase angles of spindle power peak: SO-spindle coupling was closer to synchrony in low delta power intervals with a mean SO-phase of −10.07 ± 38.71 degrees (circular mean ± SD), while high delta power intervals showed a phase shift in SO-spindle coupling with a mean SO-phase of −20.16 ± 40.65 degrees.

Interestingly, on the other hand, coupling strength analyses indicated an interrelation in the opposite direction: coupling strength for SO-spindle coupling was lower in low as compared to high delta power segments in all three naps (beta: −.057, 95% CI [−.079, −.036], t(144.9) = −5.166, p < .001, R^2^ = .114; **Fig. 2C**). Additionally, we compared SO/delta nesting index between low and high delta power segments. We found lower spindle nesting to SOs in the low compared to the high delta power condition (beta: −.027, 95% CI [−.039, −.015], t(145.66) = −4.464, p < .001, R2 = .099). Number of SO- and delta-nested spindles as well as the SO/delta nesting index for intervals of low and high delta power for each nap are displayed in **Table 1**.

**Table 1.** SO- and delta-nested spindles and the SO/delta nesting index for intervals of low and high delta power. Mean (SD) values per 30-second interval are shown.

|  | Low delta power |  |  | High delta power |  |  |
| --- | --- | --- | --- | --- | --- | --- |
| Nap (mean number of intervals per nap) | T1 (49) | T2 (51) | T3 (47) | T1 (49) | T2 (50) | T3 (47) |
| SO-nested spindles | 11.7 (7.8) | 13.1 (8.4) | 10.3 (8.3) | 16.4 (10.9) | 17.5 (9.5) | 14.2 (11.7) |
| Delta-nested spindles | 83.1 (49.9) | 95.7 (49.6) | 78.1 (51.3) | 97.3 (55.8) | 100.5 (45.6) | 89.3 (47.5) |
| SO/delta nesting index | 0.14 (0.05) | 0.13 (0.03) | 0.13 (0.04) | 0.16 (0.04) | 0.18 (0.04) | 0.15 (0.06) |

### Coupling parameters were stable over multiple naps

To further examine whether coupling strength of SO-spindle and delta-spindle coupling were stable over multiple naps within individuals, ICC values were calculated. In addition, the intra-individual stability was likewise investigated for the SO/delta nesting index. ICC estimates indicated ‘moderate’ stability of delta-spindle coupling strength (.46 (95%-CI: [.22, .66]), a ‘fair’ stability of SO-spindle coupling strength (.29 (95%-CI: [−.20, .24]) and ‘fair’ stability of the SO/delta nesting index (.31 (95 % CI [.08, .54]) according to Landis and Koch (1977) benchmarks. As such delta-spindle coupling, SO-spindle coupling and the SO/delta nesting index can be considered at least fairly stable within subjects across three naps.

### Nesting index was the most predictive measure for the ability to form novel memories

As intra-individual differences in coupling strength and SO/delta nesting index could be shown to be stable across several naps, we tested for possible associations with the ability to form novel memories, while controlling for the effect of age and gender. We found a positive association between SO/delta nesting index and superior performance during encoding (r= .35, p= .046). In addition, a slightly weaker correlation between SO/delta nesting index and immediate recall was observed(r = .33, p = .06, **Fig. 3A**). SO-spindle coupling strength (encoding: r = .08, p = .66; recall: r = .28, p = .12, **Fig. 3B**) and delta-spindle coupling strength (encoding: r = .18, p = .31; recall: r = .17, p = .34, **Fig. 3C**) were only very weakly associated with memory performance. Statistical comparisons of the correlations failed to reach significance though (all p > 0.1; Eid et al., 2011; Lenhard and Lenhard, 2014).

**Figure 3.**
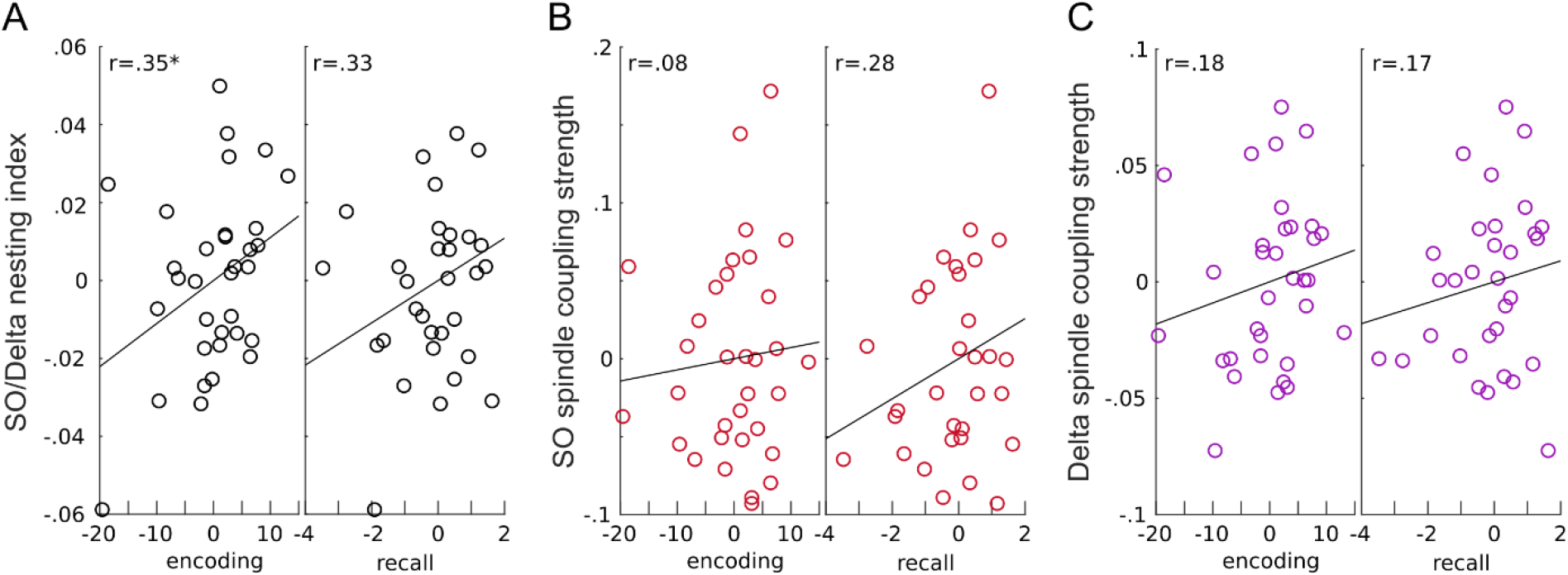
Association with memory. Scatter plots of partial correlation (after eliminating variations explained by age and gender) for VLMT encoding and immediate recall with SO/delta nesting index (A), SO-spindle coupling strength (B), and delta-spindle coupling strength (resultant vector length) (C). The SO/delta nesting index was positively associated with better encoding (p = .046) and trendwise with better immediate recall (p = .06), while only very weak correlation was found for SO and delta spindle coupling strength with memory performance. SO/delta nesting index is the ratio of SO- and delta-nested spindles. Black lines show the trend lines. \**p* < 0.05

## Discussion

Our study aimed to characterize delta waves in human NREM sleep by investigating their temporal characteristics in relation to spindles and compare their coupling to SO-spindle coupling, their intra-individual stability over multiple naps, and their correlation to the ability to form novel memories in healthy older adults. In addition, we examined the recently proposed reciprocal interaction between delta- and SO-related activities.

### Stronger SO-spindle compared to delta-spindle coupling

With regard to delta-spindle interrelations, we showed a temporal clustering of spindles in delta waves in humans, in line with an earlier finding in rats (Kim et al., 2019). We have implemented an approach that allowed us to provide detailed information about phase and strength of delta-spindle coupling, and to compare spindle coupling among oscillations of diverging frequencies such as SO and delta waves. We found similar patterns with regard to the specific timing of spindle power maxima within the slow waves’ phase: In both cases, spindle activity preferentially occurred in the late rising phase of the slow wave. However, coupling strength with spindles was stronger in SOs as compared to delta waves, which can be interpreted in line with global vs. local processes (Genzel et al., 2014). It can be assumed that SOs were higher in amplitude and more globally synchronized across cortical and sub-cortical regions (Genzel et al., 2014; Siclari et al., 2014; Bernardi et al., 2018), thereby enabling a stronger driving role on thalamic networks and on the generation as well as timed modulation of sleep spindles (Mak-McCully et al., 2017; Navarrete et al., 2020). Delta waves (Dang-Vu et al., 2008), on the other hand, with their lower amplitudes and more local patterns (Nir et al., 2011), may exhibit only moderately modulating effects on spindles, leading to lower coupling strength with thalamo-cortical spindles.

Our findings can thus be considered in line with a previous report describing how slow wave-spindle coupling is substantially influenced by the convergent effect of cortically distributed slow wave downstates (0.5 −4 Hz) on thalamic-networks (Mak-McCully et al., 2017). The authors found that a downstate in the thalamus emerges more likely when downstates occur in multiple cortical locations, probably by the abrupt silencing of depolarizing projections from the cortex. Thalamic downstates in turn trigger thalamic spindles and eventually cortical spindles. Thus, this mechanism may account for strong SO-spindle coupling. Importantly, downstates occurred about 50 % more often in the cortex than in the thalamus (Mak-McCully et al., 2017). These restricted cortical events may possibly represent delta waves. Future studies using bipolar depth recordings should further investigate distinguishing features of slow waves, namely whether only spatial distribution, or also amplitude and frequency, determine their occurrence in the thalamus.

### Interactions of delta- and SO-related activity

By exploring SO-spindle coupling and the ratio of SO-nested spindles to delta-nested spindles (SO/delta nesting index) separately in conditions of low and high delta power, we aimed to unveil the proposed competitive interactions between SOs and delta waves. Such interdependencies were reported by Kim et al. (2019), who disrupted spiking activity either during SOs or during delta waves by means of optogenetic stimulation. When delta waves were disturbed, their amplitudes were reduced, followed closely by an increase in spindle nesting to SOs. Given these findings in rats, we aimed to reveal whether interdependencies between SO and delta related activities would be similar in natural human sleep. We therefore tested whether SO-spindle coupling was more pronounced in low compared to high delta power conditions. Unexpectedly and at odds with the results of Kim and colleagues, we found higher coupling strength in high compared to low delta power condition. Surprisingly, similar to coupling strength, the SO/delta nesting index was higher in the high delta power condition. Thus, our results indeed indicate a dependency of SO-spindle coupling on delta activity in humans, but in the opposite direction as found in rats (Kim et al., 2019). Kim and colleagues proposed that cortical-subcortical structures may support reciprocal interactions between SOs and delta waves, thereby enabling a competitive balance between consolidation and homeostasis during sleep. Here, we were not able to confirm a competitive interaction between SO and delta wave activity during natural human sleep.

On the one hand, the discrepancy in findings may point to a specificity of the species and/or may arise from differences in study design. Since we were interested in the interaction between delta and SO-related activity during natural (unperturbed) sleep in humans, time periods of differing delta power were selected and compared. Thereby we were able to evaluate a temporal association between delta and SO-related activity, but could not probe causal interactions using a perturbation approach, which may account for the diverging results. Further, competitive interactions between SO and delta may also be more locally restricted and therefore only become evident in local LFP recordings but not in EEG signals providing macroscale information on neural activity (Fröhlich, 2016).

On the other hand, given that Kim and colleagues targeted firing rate during SOs or delta waves and not the oscillations themselves, changes in spindle nesting to either SO or delta waves may not have been related to modifications in slow wave activity but were in some way linked to the manipulation of local spiking activity. Indeed, previous rodent studies have shown that LFPs correlate with membrane potential oscillations of closely located neurons (Poulet and Petersen, 2008) independent of spiking activity (Okun et al., 2010). Note that while delta waves were reduced in amplitude by the optogenetic stimulation in Kim et al., SO waves were not affected, raising the question whether these findings actually reflect a direct competitive dependency between these slow waves, or whether other mediating mechanisms were involved. Thus, future studies should determine if a precisely timed manipulation of individual slow wave oscillations (e.g., by transcranial magnetic stimulation) in humans would reveal similar results as compared to those found in rats (Kim et al., 2019).

### Intra-individual temporal stability of coupling parameters and their link to memory

Several physiological sleep metrics such as time spent in different sleep stages and delta power during nocturnal sleep were shown to be stable and thus represent “trait-like” individual characteristics (Tucker et al., 2007). Cox et al. (2018) described a high intra-individual stability of SO-spindle coupling together with a high inter-individual variability, further stressing the trait-like nature of this oscillatory interaction. Our results during daytime napping were consistent with these observations; of note, delta spindle coupling strength indicated highest stability, while SO-spindle coupling strength and the SO/delta nesting index showed lower (“fair”) consistency between the three nap sessions. The lower stability of these two parameters may be due to differences between nap and nocturnal sleep. Interestingly, the SO/delta nesting index was positively associated with individual ability to form novel memories (corrected for age and gender), while SO-spindle as well as delta-spindle coupling strength by themselves were not. Assuming that delta-spindle coupling supports synaptic downscaling processes to recover the brain’s capacity to learn, we initially hypothesized the contrary relationship. In spite of the fact that this assumption could not be confirmed, the finding underscores the relevance of interactions between SO- and delta spindle nesting and points towards distinct effects of SO- and delta related activity on the ability to form novel memories. Given the exploratory nature of the present study and the fact that statistical comparisons between correlations did not reveal substantial differences, these findings have to be interpreted with caution. Future studies should disentangle in more detail how these sleep oscillations interact to impact on general memory abilities as well as sleep-dependent memory encoding and consolidation.

### Limitations

Several limitations should be considered when interpreting our findings: First, in accordance with the conventions for differentiating slow waves into SOs and delta waves (Rasch and Born, 2013; Klinzing et al., 2019) as well as previous findings that speak in favor of the frequency criterion (Achermann and Borbély, 1997; Mölle et al., 2002; Muehlroth and Werkle-Bergner, 2020), we chose to use that as the main criterion to distinguish the waveforms. In Kim et al. (2019) an amplitude criterion for the down-state was applied for differentiation into SOs and delta waves instead, implying that SOs in their study were mainly K-complexes. Our criterion does not exclude K-complexes, and since nap sleep mainly consists of N2 which contains most K-complexes, it is likely that their fraction in our SO-data is large. Future studies in both animal models and humans should investigate whether differences in differentiation criterion, or study design (nap vs. nocturnal sleep), would lead to similar results.

Second, this study was an exploratory analysis of an existing data set and thus, the design was not optimized to address the primary research questions. A) The inclusion of young subjects would lead to increased generalizability of our results (we did, however, correct for age in our analyses). B) The inclusion of nocturnal sleep, which is rich in SWS (where delta waves and SOs predominate), would be favorable.

Finally, the interval length of 30 s for analysis of SO-delta interrelations were chosen according to standard sleep scoring recommendations (Rechtschaffen and Kales, 1968; Berry et al., 2012) and seemed adequate to reach a sufficient number of slow wave events. However, no systematic investigation has been conducted on the optimal interval length for this analysis, but may possibly impact on the results.

### Conclusion

In sum, our results indicate intra-individual stability of SO-, delta- and spindle interactions, and show differences as well as dependencies between SO- and delta-related activities including spindle coupling. Further, our study suggests that SOs, delta waves and sleep spindles should be jointly considered when aiming to link sleep neurophysiology and memory formation in aging.

## Funding

This work was supported by the Deutsche Forschungsgemeinschaft (327654276 – SFB 1315, to AF and KO).

## Notes

### Competing Interest Statement

The authors have declared no competing interest.

## References

Achermann P, Borbély AA (1997) Low-frequency (< 1 Hz) oscillations in the human sleep electroencephalogram. Neuroscience 81:213–222.

Aru J, Aru J, Priesemann V, Wibral M, Lana L, Pipa G, Singer W, Vicente R (2015) Untangling cross-frequency coupling in neuroscience. Current opinion in neurobiology 31:51–61.

Baran B, Mantua J, Spencer RMC (2016) Age-related Changes in the Sleep-dependent Reorganization of Declarative Memories. Journal of Cognitive Neuroscience 28:792–802.

Berens P (2009) CircStat: A MATLAB Toolbox for Circular Statistics. 2009 31:21.

Bernardi G, Siclari F, Handjaras G, Riedner BA, Tononi G (2018) Local and Widespread Slow Waves in Stable NREM Sleep: Evidence for Distinct Regulation Mechanisms. Frontiers in human neuroscience 12:248.

Berry RB, Brooks R, Gamaldo CE, Harding SM, Marcus C, Vaughn BV (2012) The AASM manual for the scoring of sleep and associated events. Rules, Terminology and Technical Specifications, Darien, Illinois, American Academy of Sleep Medicine 176:2012.

Cox R, Mylonas DS, Manoach DS, Stickgold R (2018) Large-scale structure and individual fingerprints of locally coupled sleep oscillations. Sleep 41.

Eid M, Gollwitzer M, Schmitt M (2011) Statistik und Forschungsmethoden Lehrbuch. Beltz. In: Weinheim.

Fröhlich F (2016) Chapter 10 - LFP and EEG. In: Network Neuroscience (Fröhlich F, ed), pp 127–143. San Diego: Academic Press.

Gais S, Born J (2004) Declarative memory consolidation: mechanisms acting during human sleep. Learning & memory (Cold Spring Harbor, NY) 11:679–685.

Genzel L, Kroes MCW, Dresler M, Battaglia FP (2014) Light sleep versus slow wave sleep in memory consolidation: a question of global versus local processes? Trends in Neurosciences 37:10–19.

Göder R, Baier PC, Beith B, Baecker C, Seeck-Hirschner M, Junghanns K, Marshall L (2013) Effects of transcranial direct current stimulation during sleep on memory performance in patients with schizophrenia. Schizophrenia research 144:153–154.

Helfrich RF, Mander BA, Jagust WJ, Knight RT, Walker MP (2018) Old Brains Come Uncoupled in Sleep: Slow Wave-Spindle Synchrony, Brain Atrophy, and Forgetting. Neuron 97:221–230 e224.

Helfrich RF, Lendner JD, Mander BA, Guillen H, Paff M, Mnatsakanyan L, Vadera S, Walker MP, Lin JJ, Knight RT (2019) Bidirectional prefrontal-hippocampal dynamics organize information transfer during sleep in humans. Nature communications 10:3572.

Helmstaedter C, Lendt M, Lux S (2001) Verbaler Lern-und Merkfähigkeitstest (VLMT). Göttingen: Beltz.

Iacobucci D, Posavac SS, Kardes FR, Schneider MJ, Popovich DL (2015) The median split: Robust, refined, and revived. Journal of Consumer Psychology 25:690–704.

Kim J, Gulati T, Ganguly K (2019) Competing Roles of Slow Oscillations and Delta Waves in Memory Consolidation versus Forgetting. Cell 179:514–526 e513.

Klinzing JG, Niethard N, Born J (2019) Mechanisms of systems memory consolidation during sleep. Nat Neurosci 22:1598–1610.

Ladenbauer J, Ladenbauer J, Külzow N, Flöel A (2021) Memory-relevant nap sleep physiology in healthy and pathological aging. Sleep.

Ladenbauer J, Kulzow N, Passmann S, Antonenko D, Grittner U, Tamm S, Floel A (2016) Brain stimulation during an afternoon nap boosts slow oscillatory activity and memory consolidation in older adults. Neuroimage 142:311–323.

Ladenbauer J, Ladenbauer J, Kulzow N, de Boor R, Avramova E, Grittner U, Floel A (2017) Promoting Sleep Oscillations and Their Functional Coupling by Transcranial Stimulation Enhances Memory Consolidation in Mild Cognitive Impairment. J Neurosci 37:7111–7124.

Landis JR, Koch GG (1977) The measurement of observer agreement for categorical data. Biometrics 33:159–174.

Lemieux M, Chen J-Y, Lonjers P, Bazhenov M, Timofeev I (2014) The impact of cortical deafferentation on the neocortical slow oscillation. Journal of Neuroscience 34:5689–5703.

Lenhard W, Lenhard A (2014) Hypothesis tests for comparing correlations. Bibergau, Germany: Psychometrica.

Mak-McCully RA, Rolland M, Sargsyan A, Gonzalez C, Magnin M, Chauvel P, Rey M, Bastuji H, Halgren E (2017) Coordination of cortical and thalamic activity during non-REM sleep in humans. Nature communications 8:15499.

Maris E, Oostenveld R (2007) Nonparametric statistical testing of EEG- and MEG-data. Journal of neuroscience methods 164:177–190.

Mikutta C, Feige B, Maier JG, Hertenstein E, Holz J, Riemann D, Nissen C (2019) Phase-amplitude coupling of sleep slow oscillatory and spindle activity correlates with overnight memory consolidation. Journal of sleep research 28:e12835.

Mölle M, Marshall L, Gais S, Born J (2002) Grouping of spindle activity during slow oscillations in human non-rapid eye movement sleep. J Neurosci 22:10941–10947.

Mölle M, Marshall L, Gais S, Born J (2004) Learning increases human electroencephalographic coherence during subsequent slow sleep oscillations. Proceedings of the National Academy of Sciences of the United States of America 101:13963–13968.

Mölle M, Bergmann TO, Marshall L, Born J (2011) Fast and Slow Spindles during the Sleep Slow Oscillation: Disparate Coalescence and Engagement in Memory Processing. Sleep 34:1411–1421.

Mölle M, Eschenko O, Gais S, Sara SJ, Born J (2009) The influence of learning on sleep slow oscillations and associated spindles and ripples in humans and rats. The European journal of neuroscience 29:1071–1081.

Muehlroth BE, Werkle-Bergner M (2020) Understanding the interplay of sleep and aging: Methodological challenges. Psychophysiology 57:e13523.

Muehlroth BE, Sander MC, Fandakova Y, Grandy TH, Rasch B, Shing YL, Werkle-Bergner M (2019) Precise Slow Oscillation-Spindle Coupling Promotes Memory Consolidation in Younger and Older Adults. Scientific reports 9:1940.

Navarrete M, Schneider J, Ngo HV, Valderrama M, Casson AJ, Lewis PA (2020) Examining the optimal timing for closed-loop auditory stimulation of slow-wave sleep in young and older adults. Sleep 43.

Ngo HV, Born J (2019) Sleep and the Balance between Memory and Forgetting. Cell 179:289–291.

Nir Y, Staba RJ, Andrillon T, Vyazovskiy VV, Cirelli C, Fried I, Tononi G (2011) Regional slow waves and spindles in human sleep. Neuron 70:153–169.

Okun M, Naim A, Lampl I (2010) The Subthreshold Relation between Cortical Local Field Potential and Neuronal Firing Unveiled by Intracellular Recordings in Awake Rats. The Journal of Neuroscience 30:4440–4448.

Oostenveld R, Fries P, Maris E, Schoffelen JM (2011) FieldTrip: Open source software for advanced analysis of MEG, EEG, and invasive electrophysiological data. Computational intelligence and neuroscience 2011:156869.

Poulet JF, Petersen CC (2008) Internal brain state regulates membrane potential synchrony in barrel cortex of behaving mice. Nature 454:881–885.

Rasch B, Born J (2013) About sleep’s role in memory. Physiol Rev 93:681–766.

Rechtschaffen A, Kales A (1968) Techniques and Scoring System for Sleep Stages of Human Subjects. A Manual of Standardized Terminology.

Siclari F, Bernardi G, Riedner BA, LaRocque JJ, Benca RM, Tononi G (2014) Two distinct synchronization processes in the transition to sleep: a high-density electroencephalographic study. Sleep 37:1621–1637.

Staresina BP, Bergmann TO, Bonnefond M, van der Meij R, Jensen O, Deuker L, Elger CE, Axmacher N, Fell J (2015) Hierarchical nesting of slow oscillations, spindles and ripples in the human hippocampus during sleep. Nat Neurosci 18:1679–1686.

Tucker AM, Dinges DF, Van Dongen HP (2007) Trait interindividual differences in the sleep physiology of healthy young adults. Journal of sleep research 16:170–180.

van Dongen HPA, Vitellaro KM, Dinges DF (2005) Individual Differences in Adult Human Sleep and Wakefulness: Leitmotif for a Research Agenda. Sleep 28:479–498.

